# Mouse Genome Editing from Scratch

**DOI:** 10.1101/2023.11.20.567793

**Authors:** J.V. Popova, V.D. Bets, E.S. Omelina, L.V. Boldyreva, E.N. Kozhevnikova

## Abstract

Mouse genome modification requires costly equipment and highly skilled personnel to manipulate zygotes. A number of zygote electroporation techniques were reported to be highly efficient in gene delivery. One of these methods called i-GONAD (improved Genome-editing via Oviductal Nucleic Acids Delivery) describes electroporation-based gene transfer to zygotes *in utero*. Here we adopted this technology for mouse genome-editing from scratch minimizing the cost of equipment, operator skill and animal use. We chose the CRISPR/Cas9 system as a genome editing tool and i-GONAD as a gene delivery method to produce *IL10* gene knockout in C57BL/6 mice. Four animals out of 13 delivered pups (30.8%) were genetically compromised at *IL10* gene locus suggesting the feasibility of the approach. This report provides one of the possible technical settings for those who aim at establishing in-house mouse transgenesis pipeline at minimal cost from scratch.

**For citation:** Popova J.V., Bets V.D., Omelina E.S., Boldyreva L.V., Kozhevnikova E.N. Mouse Genome Editing from Scratch. Journal of Experimental Biology. *Year; issue (number): page. DOI*

## INTRODUCTION

Gene modification in animals includes several methodological aspects, such as anesthesia/analgesia, embryological procedures, molecular tools and delivery methods. Each of these aspects offers a choice of technical options, which vary among laboratories depending on the researchers’ preferences and skills.

Development of precise genome editing technologies increases the availability of transgenesis in laboratory animals. Many research teams may be considering in-house gene modification as an addition to their methodological repertoire. Laboratories and facilities that have long-standing experience in the production of transgenic animals tend to follow the proven techniques and may avoid technical improvements. On the contrary, newly established teams search for the optimal transgenesis methods in order to minimize the costs of equipment, reagents, and time spent to master the technology, while optimizing animal use in accordance with the 3Rs principle.

Embryo microinjection and transfer is the most laborious and skill-demanding aspect of gene modification in animals. The cost of the equipment required for this technical step makes the largest contribution to the total cost of transgenesis setup. These include a high-end microscope, micromanipulators, precise pipette pullers, and cell culture equipment, put aside outstanding operator skill for pronuclear microinjection. In recent years, embryo electroporation gained increased interest and greatly simplified gene delivery into zygotes, even though it has substantial limitations as compared to microinjection. In 2015, Takahashi and co-authors suggested gene modification by early zygote electroporation *in utero* using laboratory mice, and later rats and hamsters (Hirose *et al*., 2023; Ohtsuka *et al*., 2018; Sato *et al*., 2022; Takahashi *et al*., 2015).This method called i-GONAD (improved Genome-editing via Oviductal Nucleic Acids Delivery) shortcuts embryo collection, culture and transfer to pseudo pregnant females, simplifying gene delivery and reducing the cost of the transgenic technology. Until recently, the method was rarely reproduced by other groups, and required wider implementation into common practice. However, i-GONAD appears as an attractive alternative to conventional zygote electroporation and transfer for a newly establishing transgenesis team.

In this study, we pursued two major goals: (1) determine the minimal set of equipment, reagents and methods necessary to run gene engineering technology in mice; (2) assess the feasibility of the i-GONAD method and identify its possible shortcomings in practical use. C57BL/6 mouse strain was used here despite its limited suitability for such purposes, because we aimed to avoid backcrossing transgenic offspring to this desired genetic background. CRISPR/Cas9 (clusters of regularly interspaced short palindromic repeats) / CRISPR-associated nuclease 9) was used as a genome editing tool to introduce simple gene mutations.

The method presented here implies that the personnel is experienced with mouse handling, anesthesia/analgesia, surgical manipulations, and is supervised by a veterinarian doctor.

Our data suggest that CRISPR/Cas9-assisted genome editing combined with i-GONAD delivery method is suitable to introduce gene knockouts in C57BL/6 mice by unexperienced users. The approach described here might represent the simplest and cheapest set of technologies to establish a mouse gene modification method from scratch (Figure 1). However, i-GONAD was associated with significant embryo toxicity and high newborn death rate, which should be considered while implementing this technique.

**Figure 1.**
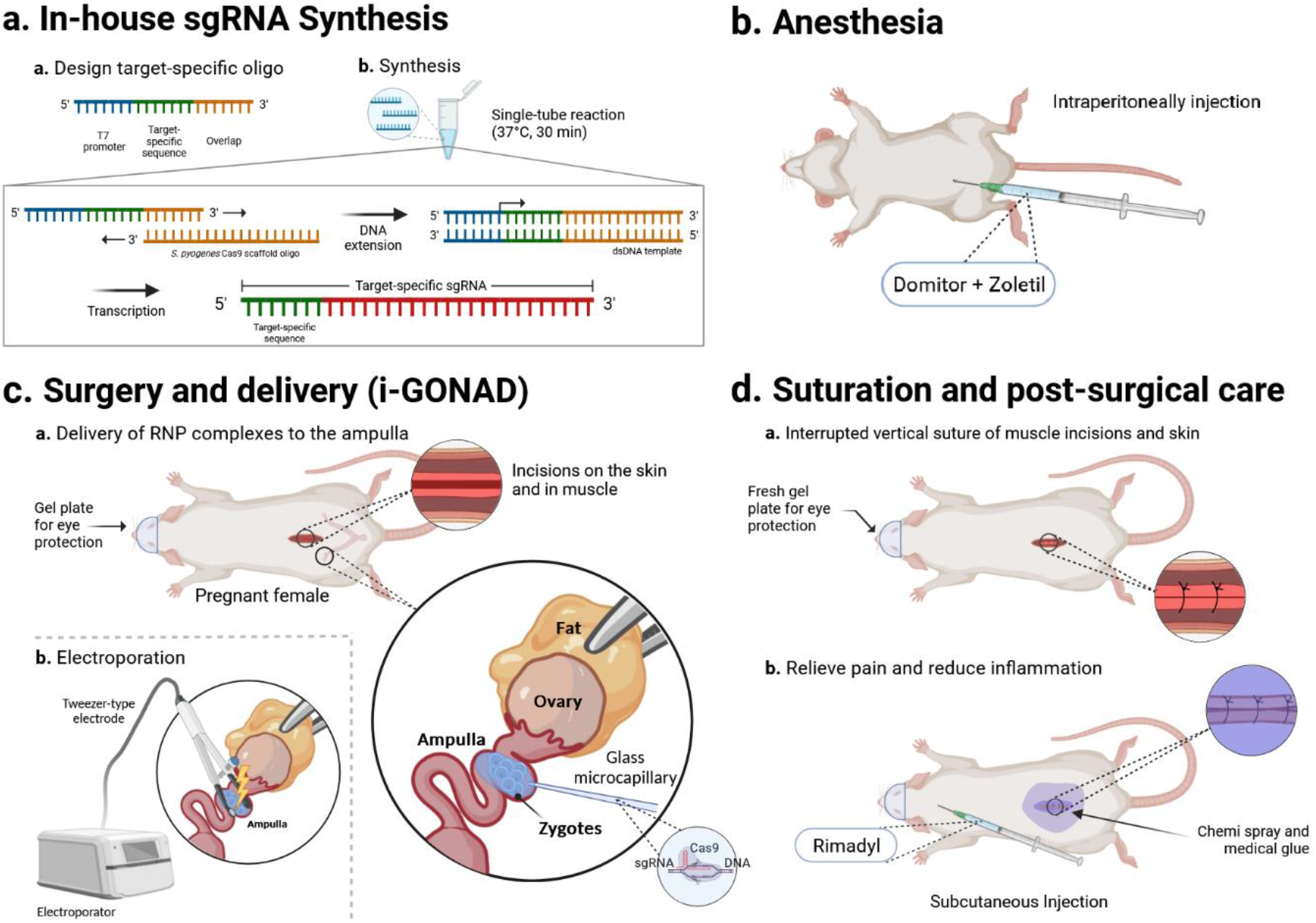
Scheme of CRISPR/Cas9 genome editing with intra-oviduct injection. **A.** In-house sgRNA synthesis. **B.** Anesthesia. **C.** i-GONAD (improved Genome-editing via Oviductal Nucleic Acids Delivery) method. **D.** Suturation and post-surgical procedures. This figure was created with biorender.com (accessed on 05 October 2023).

## MATERIALS

### Animals

The experiments were performed in the experimental animal sector at Scientific Research Institute of Neurosciences and Medicine (SRINM) (Novosibirsk, Russia). The study was conducted using inbred strain C57BL/6JNskrc (our in-house C57BL/6J sub-colony): female (n=40; 8-14 weeks old) and male (n=8; 6-20 weeks old) mice. Zygote survival and electroporation efficiency were performed using outbred strain CD-1: female (n=20; 8-14 weeks old) and male (n=8; 6-20 weeks old) mice.

Adult female mice were housed in groups of 4-5 animals per cage, adult male mice were housed individually. All animals were housed in open cages in 12 h/12 h light/dark photoperiod (00:00-12:00-00:00) under standard conditions. Food and water were provided *ad libitum*. The cages of pregnant females were not changed for two weeks, starting 7 days before delivery. Pregnant and lactating females were fed protein-and fat-rich food pellets (dry dog food for adult small breed “Opti Balance” with chicken, Purina Pro Plan, USA) in addition to the regular chow.

### Ethical statement

All procedures were conducted under Russian legislation according to the standards of Good Laboratory Practice (directive #267 from 19.06.2003 of the Ministry of Health of the Russian Federation), institutional bioethical committee guidelines and the European Convention for the protection of vertebrate animals. All procedures were approved by the Bioethical committee at SRINM, protocol #5 dated 16.02.2023. All results are reported consistently with ARRIVE guidelines (Du Sert *et al*., 2020).

### Reagents, equipment

All reagents and equipment are described in detail in Supplementary Materials in a protocol format.

### Software

FinchTV was used to analyze Sanger DNA Sequencing data (Geospiza Inc., USA). CHOPCHOP web tool was used to design guide RNA (gRNA) target sequences (chopchop.cbu.uib.no; accessed on 20 September 2023). STATISTICA12 software was used to calculate statistical significance.

## METHODS

### Preparation of genome editing mixture

The single guide RNAs (sgRNAs) for the mouse *IL10* gene were synthesized using target-specific oligo design EnGen sgRNA Synthesis Kit (NEB, cat. #E3322S) according to the protocol provided by the manufacturer (neb.com/en/protocols/2016/05/11/engen-sqrna-synthesis-kit-s-pyogenes-protocol-e3322; accessed on 20 September 2023). As a result, the following nucleotides were used for sgRNA synthesis: IL10_ex1_sg (for the *IL10* exon 1) and IL10_ex2_sg (for the *IL10* exon 2) (oligonucleotide sequences can be found in Supplementary Materials). Synthesized sgRNAs were purified using Monarch RNA Cleanup Kit (NEB, cat. #T2040S) following manufacturer’s instructions. Concentration of pure sgRNAs was detected using NanoDrop.

To prepare a genome editing solution, 30 µM of each sgRNAs and 1 mg/ml for the Cas9-NLS protein (NEB, cat. #M0646T) were diluted in Opti-MEM I Reduced Serum Medium (ThermoFisher Scientific, cat. #31985062) and used as 1.5 µl/oviduct for *in utero* electroporation.

### Plasmid cloning and sgRNA *in vitro* testing

*IL10* target loci were cloned in a plasmid vector to test the efficiency of the designed sgRNAs *in vitro*. Loci-specific primer sets IL10_ex1_gDNA_F / IL10_ex1_gDNA_R and IL10_ex2_F / IL10_ex2_gDNA_R (oligonucleotide sequences can be found in Supplementary Materials) were used for PCR amplification of genomic DNA in a total 25 µl reaction volume. The PCR products were gel-purified using DNA and RNA agarose gel isolation kit (Biolabmix, cat. #N-Gel-250) and ligated with the pBlueScript SK (+) vector +4°C overnight using T4 DNA ligase (Evrogen, cat. #LK001). The vector was previously digested at 37°C for 1 h with EcoRV endonuclease (SibEnzyme, cat. #SE-E059). 1 µl of the ligation mixture was used to transform *E. coli* TOP 10 electrocompetent cells. The final constructs, containing fragments of the *IL10* exon 1 and exon 2, were verified by Sanger sequencing. The plasmids were further linearized with ScaI endonuclease (New England Biolabs Inc., cat. #R3122) at 37°C for 1 h before being used for Cas9 digestion. *In vitro* cleavage reactions were performed in a ratio of 20:120:1 (Cas9-NLS:sgRNA:linearized DNA construct) in a total volume 30 µl at 37°C for 1 h following manufacturer’s instructions (NEB, cat. #M0646). Each reaction was mixed with loading dye and resolved in 1% agarose gel in the TAE (Tris base/Acetic acid/EDTA) buffer.

### *In utero* experiment (i-GONAD)

The step-by-step procedure of CRISPR/Cas9 genome-editing with intra-oviduct injection (anesthesia, analgesia, *in utero* injection and electroporation, mouse surgery and post-surgical care) are described in full detail in Supplementary Materials.

### Testing electroporation procedure in embryos

To determine the damage of the electroporation procedure and the potential embryotoxicity from *IL-10* construct, we assessed zygote survival in vitro. Test CD-1 animals were electroporated *in utero* using *IL10* sgRNA and Cas9 in Opti-MEM I Reduced Serum Medium (ThermoFisher Scientific, cat. #31985062) and control CD-1 mice were electroporated with 1× PBS in Opti-MEM medium. All manipulations were performed in anesthetized CD-1 pregnant females (0.7 d.p.c.) without hormonal stimulation. In both experimental groups, the solution was introduced into the ampulla and then electroporated (the detailed procedure is described in Supplementary Materials). Right after the electroporation, zygotes were washed out of the ampulla into KSOM Embryo Medium (Sigma, cat. #MR- 101-D) with hyaluronidase (Roanal, cat. #08091) (10 mg/ml in KSOM medium) onto a culture plate covered with mineral oil (Sigma-Aldrich, cat. #M8410-100ML). Then the zygotes were incubated at 37°C in 5% CO_2_ for 24 h, after which the number of dividing cells was counted.

To confirm electroporation efficiency, 4kDa FITC-dextran (Sigma-Aldrich, cat. #FD4-1G) (10 µg/µl in Opti-MEM medium) was injected into the ampulla in the volume of 1.5 µl/oviduct followed by electroporation. Zygotes were washed out and analyzed using fluorescent microscope.

### Offspring genotyping

At weaning, the offspring were labeled on the ears and these tissue samples were used to purify genomic DNA and PCR-genotype the animals. The detailed procedure of DNA purification is described in Supplementary Materials. Briefly, ear samples were lysed and DNA was purified using chloroform fractionation and ethanol precipitation. IL10_ex1_gDNA_F / IL10_ex1_gDNA_R and IL10_ex2_F / IL10_ex2_gDNA_R primer pairs were used for genotyping (oligonucleotide sequences can be found in Supplementary Materials). Genomic DNA samples were amplified using BioMaster HS-Taq PCR-Color (2×) (Biolabmix, cat. #MHC010-200) according to the manufacturer’s recommendations. DNA samples from intact C57BL/6 animals were used as a positive control and mQ H_2_O ‒ as a negative control. The PCR samples were further analyzed in a 2% agarose gel in 0.5× TBE (Tris/Borate/EDTA) buffer and isolated from gel using DNA and RNA agarose gel isolation kit (Biolabmix, cat. #N-Gel-250). The resulting DNA samples were used for Sanger sequencing using BigDye Terminator v3.1 Cycle Sequencing Kit (ThermoScientific, cat. #4337455).

## RESULTS

### *In vitro* cleavage assay

To analyze the *in vitro* efficiency of the synthesized sgRNAs, we performed *in vitro* Cas9 nuclease assay. The plasmids containing murine *IL10* target genome loci were cleaved with ScaI endonuclease, which linearized the vectors. After that, the Cas9 ribonuclear (RNP) complex cleaved the vectors resulting in two linear fragments. We assume that ScaI efficiency is close to 100%, thus the unhydrolyzed fragments result from incomplete cleavage by the RNP complex. The cleavage rate was higher for exon 1-targeting IL10_ex1_sg, as compared to exon 2-targeting IL10_ex2_sg. We used both sgRNAs in one RNP mixture for further procedures (Figure 2A).

**Figure 2.**
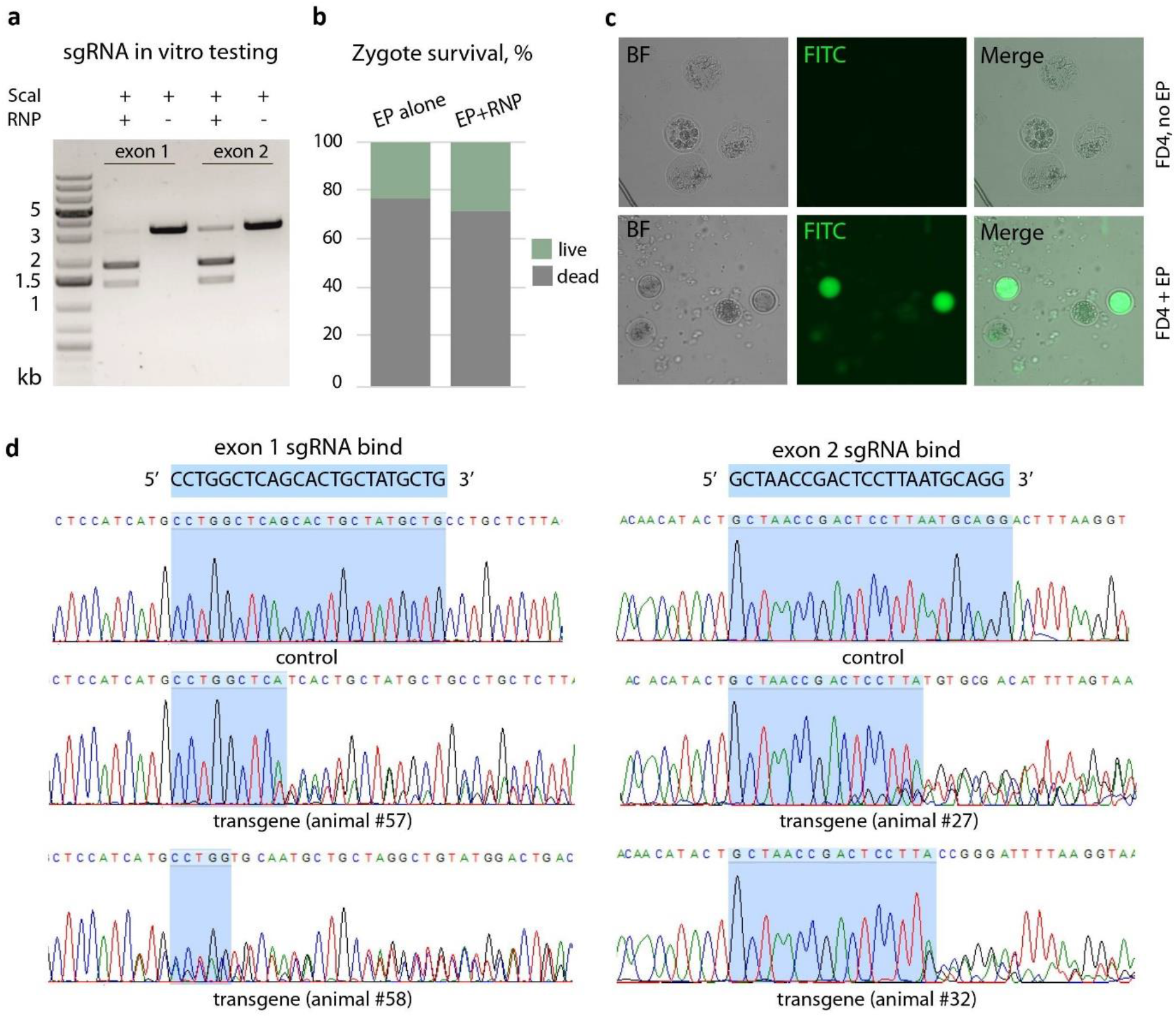
i-GONAD adaptation and testing. **A.** sgRNA (single guide RNA) *in vitro* testing. Agarose gel electrophoresis of ScaI-linearized pBlueScript SK (+) vector carrying a fragment of either *IL10* exon 1 or exon 2 genomic region. RNP – ribonuclear complexes of sgRNA and Cas9 protein. **B.** Zygote survival *in vitro* after *in utero* electroporation. EP – electroporation. **C.** *In utero* electroporation efficiency test using 4kDa FITC-dextran (FD4). **D.** Sanger sequencing of *IL10* genomic regions in F1 progeny.

### Embryo survival after electroporation

We next tested the effect of electroporation on embryo survival *in vitro* and evaluated the potential toxic effect of in-house synthesized sgRNA. We electroporated the oviducts of 0.7 d.p.c CD-1 pregnant females with either RNP in Opti-MEM or PBS in Opti-MEM, washed the embryos and kept them in cell culture conditions overnight. The following day we found that of about 100 living zygotes, 22.76% were dividing in the control group and 28.74% ‒ in the RNP group, indicating that RNP complexes had no additional toxic effect on embryos, but electroporation itself resulted in about 70% mortality in zygotes (Figure 2B).

We further tested whether electroporation conditions used as published previously resulted in gene delivery to the embryos using FITC-dextran as a fluorescent substitute for nucleic acids (Sato *et al*., 2020). We found that indeed electroporation resulted in a strong FITC signal in about 40% of electroporated zygotes, whereas no signal was observed in non-electroporated control (Figure 2C). This result was consistent with previously reported data that gene delivery via *in utero* electroporation is not 100% efficient (Garcia-Frigola *et al*., 2007; Shinmyo *et al*., 2016).

### Gene modification using i-GONAD

We next applied *IL10* RNP mixtures to zygotes of C57BL/6 mice *in utero* using i-GONAD approach. In a total of 7 attempts we used 40 female mice aged 8-14 weeks and 8 sexually experienced male mice aged 6-20 weeks. All C57BL/6 females were superovulated for i-GONAD (see Supplementary Protocols) before mating. Of 40 females only 30% gave birth, reflecting high mortality rate in embryos after electroporation as we observed in the above *in vitro* testing. These 40 females gave birth to 58 pups, of which only 13 (22.4%) survived as a result of cannibalism from mothers (Table 1). Of the remaining 13 pups, 4 pups (30.8%) were compromised in the *IL10* genomic region corresponding to the sgRNA binding site and downstream of it. Two pups were compromised in exon 1 and two other pups were compromised in exon 2 of the *IL10* gene (Figure 2D), however no genomic deletions were found in this experiment.

**Table 1.** Delivery rate, pup survival and efficiency of transgenesis.

| Experiment No | Number of females mice | Females gave birth | Pups were born | Survived pups | Pups with compromised *IL10* genomic region |
| --- | --- | --- | --- | --- | --- |
| 1 | 5 | 0 | 0 | 0 |  |
| 2 | 5 | 0 | 0 | 0 |  |
| 3 | 5 | 1 | 3 | 3 |  |
| 4 | 5 | 3 | 22 | 0 |  |
| 5 | 10 | 5 | 26 | 7 | 2 (in exon 2) |
| 6 | 7 | 2 | 4 | 0 |  |
| 7 | 3 | 1 | 3 | 3 | 2 (in exon 1) |
| **Total** | **40** | **12**<br>**(30%)** | **58** | **13**<br>**(22.4%)** | **4**<br>**(30.8% of**<br>**survived pups)** |

## DISCUSSION

In-house production of transgenic animals can be beneficial for groups that specialize in animal genetics and physiology and those aiming to obtain unique strains, for example, personalized animal models. However, this is only possible if the genome editing technology is simple enough to be implemented into the daily routine of the average animal laboratory. In addition, affordable mouse transgenesis should not require a long list of additional costly equipment and fit a laboratory’s budget.

CRISPR/Cas9 technology has significantly improved transgenic animal production, as it increased the accuracy of gene editing and bypassed the need for embryonic stem cells (Jon Cohen, 2016). At the same time, the technology of transgenesis is based on the labor-intensive stage of introducing molecular mixtures into the pronucleus of zygotes, which requires outstanding employee skills and expensive equipment. At the moment, this delivery method is being actively replaced by electroporation of zygotes, which greatly simplifies the transgenesis method (Modzelewski *et al*., 2018). However, this approach still sacrifices female mice for zygote production, requires an additional group of vasectomized males to mate pseudo pregnant recipient females, and necessitates embryos to be cultured *in vitro*. These aspects introduce additional costs and complexity that limit widespread adoption of mouse genome editing.

The i-GONAD method used here avoids the above limitations and makes transgenesis accessible virtually to any laboratory that already works with mice. This method was proven to be effective in several animal species and different mouse strains (Hirose *et al*., 2023; Ohtsuka *et al*., 2018; Sato *et al*., 2022; Takahashi *et al*., 2015). In our hands, standard published electroporation parameters appeared toxic to embryos (Ohtsuka *et al*., 2018). Imai and colleagues suggest using 3 transfer pulses instead of 6 suggested originally to increase embryo survival in different mouse strains, however, this approach resulted in no genome modifications for C57BL/6 mice (Imai *et al*., 2022).

High levels of cannibalism toward newborn pups is another technical caveat that we encountered in this study. Cannibalism is common in the widely used inbred strains such as C57BL/6 and BALB/c, while for some genetically modified strains, this risk is even higher (Carter *et al*., 2002; Weber *et al*., 2013). We observed an unusually high level of cannibalism among test females so that about 75% of pups did not survive the first days of life, which agrees with the data published previously (Carter *et al*., 2002). While this report was prepared, Melo-Silva and co-authors published their results on adaptation of the i-GONAD method in C57BL/6, where they describe similar challenges (Melo-Silva *et al*., 2023). The authors suggest co-housing synchronized pregnant C57BL/6 mice with companion pregnant FVB/NJ females, so that the later assist in pup survival (Melo-Silva *et al*., 2023). This could be a great addition to the presented here setup as it might significantly improve i-GONAD efficiency.

Overall, the approach suggested here is feasible to provide an effective and relatively simple start for the laboratory that had no previous experience with mouse embryology or transgenesis.

## Acknowledgements

We kindly thank the organizers and speakers of the EMBO Practical Course 2019 on mouse genome engineering held in Max Planck Institute of Molecular Cell Biology and Genetics (Dresden, Germany) for sharing their expertise in zygote electroporation and mouse genome editing.

## Ethics

The study was approved by the Bioethical committee of the Scientific Research Institute of Neurosciences and Medicine, protocol #5 dated 16.02.2023.

## MATERIALS

Listed below are reagents, equipment and devices that are used in this study. Comparable materials from other vendors may also be used in place of these.

### Reagents

#### Reagents for preparation of genome editing mixture and in vitro cleavage assay

1. EnGen sgRNA Synthesis Kit (NEB, cat. #E3322S)
2. Target specific DNA oligonucleotides
3. Monarch RNA Cleanup Kit (NEB, cat. #T2040S)
4. EnGen Spy Cas9-NLS protein (NEB, cat. #M0646T)
5. GeneRuler 1 kb Plus DNA Ladder (Thermo Fisher Scientific, cat. #SM1332)
6. T4 DNA ligase (Evrogen, cat. #LK001)
7. EcoRV endonuclease (Sibenzyme, cat. #SE-E059)
8. ScaI endonuclease (New England Biolabs Inc., cat. #R3122)
9. pBlueScript SK (+) vector
10. *E. coli* TOP10 electrocompetent cells
11. Nuclease-free water

#### Reagents for i-GONAD

1. EmbryoMax Advanced KSOM Embryo Medium (Sigma, cat. #MR-101-D)
2. Mineral oil (Sigma-Aldrich, cat. #M8410-100ML)
3. Opti-MEM I Reduced Serum Medium (ThermoFisher Scientific, cat. #31985062)
4. Hyaluronidase (Roanal, cat. #08091)
5. Pregnant Mare Serum Gonadotropin (PMSG) (Prospec, cat. #HOR-272)
6. Human Chorionic Gonadotropin (HCG) (Prospec, cat. #HOR-250)
7. Commercial drug “Zoletil 100” (aqueous solution medetomidine hydrochloride, 100 mg/ml) (for veterinary medicine) (Virbac, France)
8. Commercial drug “Domitor” (aqueous solution of medetomidine hydrochloride, 1 mg/ml) (for veterinary medicine) (Orion Pharma, Finland)
9. Commercial drug “Rimadyl” (solution for injections, 5%) (for veterinary medicine) (Zoetis, Brazil)
10. Fluorescein isothiocyanate-dextran (FITC-dextran) (Sigma-Aldrich, cat. #FD4-1G)
11. Opti-MEM medium (Thermo Fisher Scientific, cat. #31985062)
12. Trypan Blue Stain, 0.4% (Thermo Fisher Scientific, cat. #T10282);
13. Phosphate buffer saline (PBS) without Ca2+ and Mg2+, pH 7.2 (Thermo Fisher Scientific, cat. #70013032)
14. Normal saline (solvent for the preparation of dosage forms for injection), 0.9%
15. Medical glue “BF-6”, alcohol solution for external use (Verteks, Russia)
16. Chemi spray, anti-inflammatory and antibacterial drug (for veterinary medicine) (Industrial Veternaria S.A. Invesa, Spain)
17. Agarose (Bioron, cat. #604005)
18. Disinfectant “Kombidez” (KiiltoClean, Russia)
19. Ethahol, 70% *(for disinfection)*
20. Iodine alcohol solution, 5%
21. mQ H_2_O

#### Reagents for genotyping

1. BioMaster HS-Taq PCR-Color (2×) (Biolabmix, cat. #MHC010-200)
2. DNA and RNA agarose gels isolation kit (Biolabmix, cat. #N-Gel-250)
3. BigDye Terminator v3.1 cycle Sequencing kit (ThermoScientific, cat. #4337455)
4. Proteinase K (20 mg/ml) (Sibenzyme, cat. #E347)
5. SDS (Applichem, cat. #A1112.0500)
6. EDTA (0.5M, pH=8.0) (Sigma, cat. #BCBQ4662V)
7. Agarose (Bioron, cat. #604005)
8. TrisHCl (1M, pH=8.0)
9. NaCl (5M)
10. KAc (5M)
11. Chloroform
12. Ethanol, 96%
13. mQ H_2_O

### Equipment

#### Equipment used in all procedures

1. 1.5 ml microcentrifuge tube (SSI, cat. #EP-150-J)
2. Falcon: 15 and 50 ml (Axygen, cat. #SCT-15ml-500, cat. #SCT-50ml-500)
3. Pipets tips with filters: 10, 200, and 1000 µl (Accumax, cat. #Ac-AT-10-S-F-R; NEST, cat. #N-312012; SSI, cat. #SSI-4337NSFS)
4. Automatic pipettes: 0.5-10, 2-20, 20-200, and 100-1000 µl (IKA Pette vario, cat. #0020011211, #0020011213, #0020011215, #0020011216)
5. Latex gloves: S, M, and L size (Kimberly-Clark, cat. #57371, #57372, #57373)
6. Glass bottle: 100, 250, 500, and 1000 ml (Rasotherm, cat. #95206001, #95206002, #95206003, #95206004)
7. Parafilm M (Pechiney Plastic Packaging Company, cat. #PM 996)

#### Equipment for i-GONAD

1. Clothes for surgery: sterile masks, surgical caps, shoe covers, and medical gowns or suits
2. Petri dishes: 3.5 and 10 cm diameter (Biologix, cat. #07-3035; Next, cat. #704002)
3. Insulin syringes for single use, three-component with fixed ultra-micro needle 29G (0.33×12.7 mm), 1 ml (SFM, cat. #U-100)
4. Syringes for single use, three-component, needle 21G (0.3×40 mm), 5 ml (INEKTA, cat. #1016542)
5. Syringes, Luer Lock, for single use, three-component, without needle, 50 ml (Vogt Medical GmbH, cat. #0154-375-291-137)
6. Syringe filter, 0.22 µm (TTP, cat. #99722)
7. Surgical PGA (polyglycolide or polyglycolic acid) absorbable suture thread with needle, USP 5/0, 75 cm, HR-20 (Lintex, cat. #HR-20)
8. Painting tape *(for fixing the paws of the animal)*
9. Surgical cotton wool
10. Sterile, absorbent wipes, density 50 g/m^2^, 10×10 cm (Mastmed, Russia, cat. #613013)
11. Lint-free wipes for cleaning optics, 21.3×11.4 cm, (Kimtech Science, cat. #7552)
12. Paper wipes
13. Filter paper
14. Silicone mat (size 30×30 cm)
15. Mouse weighing container
16. Protective shield or goggles *(necessary to protect the face during the haircut of the animal!)*
17. Mouth pipette (mouth piece, 20 or 200 µl pipette tip with filter, silicon tubing ∼30 cm, 2 pcs. 200 µl pipette tips) with glass micropipettes for intra-oviduct injection
18. Glass capillary, 1.0 mm O.D. × 0.58 mm I.D. (Harvard Apparatus Ltd, cat. #GC100F-10)
19. PE package for the collection and disposal of class B medical waste, 5 L, 33×30 cm

#### Devices

1. Electroporator (*in vivo* and *in vitro* electroporator CUY21EDIT II) (BEX CO LTD, cat. #CUY21EDIT2)
2. Tweezer-type electrode (BEX CO LTD, cat. #LF650P3)
3. Mini vortex centrifuge “Micro-spin” (Biosan, cat. #BS-010201-AAA)
4. Centrifuge (Eppendorf, cat. #5417R)
5. T100 Thermal Cycler (Bio-rad, cat. #1861096)
6. Thermostat (Biosan, cat. #BS-010401-QAA)
7. NanoDrop (NanoDrop Technologies, cat. #ND-1000)
8. Microscope (Zeiss, cat. #Stemi 2000)
9. Vertical needle puller (micro-forge) for obtaining micropipettes from glass capillaries by pulling the tips
10. Thermal mats (for reptiles) (max temperature 38°C): size 14×15 cm (1 pc.) and 28×53 cm (2-3 pcs.)
11. Laboratory scales (OHAUS corp., cat. #Scout II/SC2020)
12. Trimmer (hair clipper)
13. Stopwatch timer

#### Surgical instruments

1. Scalpel Blades (RWD, cat. #S31010-01)
2. Scalpel Handles (RWD, cat. #S32004-13)
3. Scalpel Blade Remover (RWD, cat. #S33009-06)
4. Micro Forceps-Str, head length 34 mm, tip 0.35×0.55 mm, 11 cm (RWD, cat. #F13002-11)
5. Dressing Forceps-Str, tip width 1.4 mm, teeth length 19 mm, 15.5 cm (RWD, cat. #F12016-15)
6. DIEFFENBACH Bulldog Clips-Str / 14×4 mm / 48 mm (RWD, cat. #R33001-48)
7. MAYO-HEGAR Needle Holders-Str / 17.5×1.75 mm / 14 cm (RWD, cat. #F31034-14)
8. IRIS T.C. Scissors (Round Type)-S/S Str / 9 cm (RWD, cat. #S18004-09)
9. Dressing Forceps w/o Serrations-45°Cvd, S/S, Tip 0.5×0.2 mm, 10 cm (RWD, cat. #F12013-10)
10. VANNAS Spring Scissors (Triangular)-S/S Str / 8×1.55 mm / 8 cm (RWD, cat. #S11001-08)
11. VANNAS Spring Scissors (Triangular)-S/S Str / 3.5×1.3 mm / 8 cm (RWD, cat. #S11035-08)

## METHODS

### MOLECULAR WORK

#### Preparation of genome editing mixture

The sgRNA for the *IL10* gene was designed using CHOPCHOP (chopchop.cbu.uib.no; accessed on 20 September 2023) web tool for selecting target sites for CRISPR/Cas9 and target-specific oligo design service from EnGen sgRNA Synthesis Kit, *S. pyogenes* Protocol (NEB, cat. #E3322). As a result, we used two DNA oligonucleotides ‒ IL10_ex1_sg (for the *IL10* exon 1) and IL10_ex2_sg (for the *IL10* exon 2). Oligonucleotide sequences:

IL10_ex1_sg – 5’ TTCTAATACGACTCACTATAGCAGCATAGCAGTGCTGAGCCGTTTTAGAGCTAGA 3’

IL10_ex2_sg – 5’ TTCTAATACGACTCACTATAGCTAACCGACTCCTTAATGC GTTTTAGAGCTAGA 3’

The sgRNAs for exon 1 and exon 2 of the *IL10* gene were synthesized using the primers IL10_ex1_sg and IL10_ex2_sg, respectively, and the EnGen sgRNA Synthesis Kit (NEB, cat. #E3322S). Synthesized sgRNAs were purified using Monarch RNA Cleanup Kit (NEB, cat. #T2040S) following manufacturer’s instructions. Concentration of pure sgRNAs was detected using NanoDrop.

To prepare a genome editing solution, CRISPR components were mixed together, so that the final concentration of components was 1 mg/ml for the Cas9-NLS protein (NEB, cat. #M0646T) and 30 µM for sgRNAs. This solution was diluted using Opti-MEM I Reduced Serum Medium (ThermoFisher Scientific, cat. #31985062) to adjust the volume to 1.5 µl/oviduct during electroporation.

### Plasmid cloning and sgRNA *in vitro* testing

To analyze synthesized sgRNAs, we performed *in vitro* Cas9 nuclease assay. To do that, the PCR amplification of target loci *IL10* was performed in a total 25 µl solution using the primer sets IL10_ex1_gDNA_F / IL10_ex1_gDNA_R and IL10_ex2_gDNA_F / IL10_ex2_gDNA_R for the regions of the *IL10* exon 1 and exon 2, respectively. Oligonucleotide sequences:

IL10_ex1_gDNA_F – 5’ ATTGCATGGTTTAGAAGAGGGA 3’

IL10_ex1_gDNA_R – 5’ TTATTGTCTTCCCGGCTGTACT 3’

IL10_ex2_gDNA_F – 5’ AGTCCTTGCATTCACGTTCTTT 3’

IL10_ex2_gDNA_R – 5’ GACTTACTGGAATGGTGATCTGTTGC 3’.

The PCR products were gel-purified using DNA and RNA agarose gels isolation kit (Biolabmix, cat. #N-Gel-250). After purification the PCR products and the pBlueScript SK (+), previously digested with EcoRV endonuclease (Sibenzyme, cat. #SE-E059) at 37°C for 1 h, were ligated at +4°C overnight with T4 DNA ligase (Evrogen, cat. #LK001). Then, 1 µl of the ligation mixture was used to transform *E. coli* TOP10 electrocompetent cells. The final constructs, containing fragments of the *IL10* exon 1 and exon 2, were verified by Sanger sequencing and digested with ScaI endonuclease (New England Biolabs Inc., cat. #R3122) at 37°C for 1 h to linearize the plasmids.

*In vitro* cleavage reactions were performed with a ratio of 20:120:1 (Cas9-NLS:sgRNA:DNA construct) in a total volume 30 µl at 37°C for 1 h following manufacturer’s instructions (NEB protocol cat. #M0646). Each reaction was mixed with loading dye and resolved in 1% agarose gel in the 1× TAE (Tris base/Acetic acid/EDTA) buffer.

#### WORK WITH ANIMALS

*Note: all manipulations with animals are performed in special clothing (sterile masks, surgical caps, shoe covers, and medical gowns or suits)!*

### Preparation of pregnant female mice (hormone stimulation)

We use superovulation in C57BL/6 female mice, since it was shown previously that genetic modification in inbred mouse strains of interest such as C57BL/6 is still challenging because of their low fertility and embryo survival. (Melo-Silva *et al*., 2023) The i-GONAD procedure should be carried out on day 0.7 after fertilization (d.p.c., days past coating) (Ohtsuka *et al*., 2018), thus we used the following protocol for mating females with males adapted to the light regime of our animal facility (12 h/12 h light/dark photoperiod, [00:00- 12:00-00:00]):

1. Day 1, 11:00 a.m. With hands (in gloves) carefully remove the animal from the cage and fix it in the left hand: the thumb and forefinger fix the scruff of the animal, and the little finger presses the tail to the palm of your hand. Intraperitoneally inject PMSG (5 IU) (Prospec, cat. #HOR-272) in the amount 100 μl/mouse using insulin syringes. PMSG solution is prepared in accordance with the manufacturer’s instructions, then divided to 1 ml aliquots and stored at -20°C. *Note: Do not filter hormone solutions!*
2. Day 2, pause.
3. Day 3, 11:00 a.m. Take the animal in your hands, as described in paragraph #1. Intraperitoneally inject HCG (5 IU) (Prospec, cat. #HOR-250) in the amount 100 μl/mouse using insulin syringes. HCG working solution is prepared in accordance with the manufacturer’s instructions, then divided to 1 ml aliquots and stored at -20°C. *Note: Do not filter hormone solutions!*
4. After the injection of HCG directly put females in male’s cage (1-2 females per 1 male).
5. Day 4, 10.00 a.m. Check females for the presence of vaginal plugs (a sign of successful mating). Further manipulations with females should be performed starting from 12:30 a.m. (approximately 0.7 d.p.c.).

For CD-1 female mice, no hormonal stimulation was applied.

### Pre-surgical preparation

#### Preparation of glass micropipettes for intra-oviduct injection

The preparation of capillaries was carried out according to the protocols published previously (Meyer-Dilhet and Courchet, 2020). Using a vertical needle puller (micro-forge), make micropipettes from glass capillaries by pulling the tips. The glass capillary should be thin and sharp enough not to damage the tissues during the introduction of solutions into the oviduct. And, at the same time, its inner diameter at the tip must be sufficient to easily blow out the solution using the "Mouth pipette" system (Figure S1.A). One experiment (4-5 animals) requires 3-4 capillaries.

#### Workspace preparation

Prepare workspace for surgery (intra-oviduct injection) (Figure S1.B-D).

1. Place a clean mouse cage on thermal mat sized 28×53 cm (set the heating temperature to 35-37°С) using no more than 4-5 animals per cage.
2. Place sterile cotton balls in a clean glass bottle (volume 250 or 500 ml) in an iodine alcohol solution (5%).
3. Prepare a covering material for the surgical field. In absorbent wipes (10×10 cm) (Mastmed, Russia, cat. #613013) make a 2×2 cm hole in the center of each wipe. Before surgery, sterilize the wipes.
4. Prepare filter wipes to remove blood and other liquids during surgery. Cut filter paper about 1×1 cm in size, sterilize before use in a separate container. During surgery, transfer part of the sterile filter wipes from the sterilization container to a sterile Petri dish.
5. Prepare covering wipes for the oviduct during electroporation. Cut wipes for optics (Kimtech Science, cat. #7552) into strips about 3×7 mm in size, then place these pieces in a separate container (preferably a small air-tight glass cup) and autoclave.
6. Cut the painting tape into strips of a size convenient for fixing the animal’s paws during the surgery operation.
7. Prepare a fresh solution of 1× PBS (Thermo Fisher Scientific, cat. #70013032), filter using syringe filter (TTP, cat. #99722) and place 2-3 ml into a sterile Petri dish (diameter 3.5 cm). It is needed for wetting filter paper wipes, covering wipes for the oviduct and electrodes.
8. Place sterile instruments and surgical PGA (polyglycolide or polyglycolic acid) absorbable suture with needle into a sterile Petri dish (diameter 10 cm).
9. Prepare the workspace for surgery. Cover the silicone mat (size 30×30 cm) with a sheet of filter paper (size ∼25×25 cm) and fix it with painting tape. Place a thermal mat (size 14×15 cm) on top of the filter paper and set the heating temperature to 35-37°С.
10. Prepare the genome editing solution (the final concentration of the Cas9-NLS protein and sgRNAs were 1 mg/ml and 30 µM, respectively) and add Trypan Blue Stain (0.4%) (Thermo Fisher Scientific, cat. #T10282), required to visualize the injection of the solution into the oviduct (10 parts injection solution to 1 part dye). One mouse requires 3 µl of the prepared solution (1.5 µl per oviduct). Then, using the assembled "Mouth pipette" system, draw the required amount of solution into the ready-made capillary.

**Figure S1.**
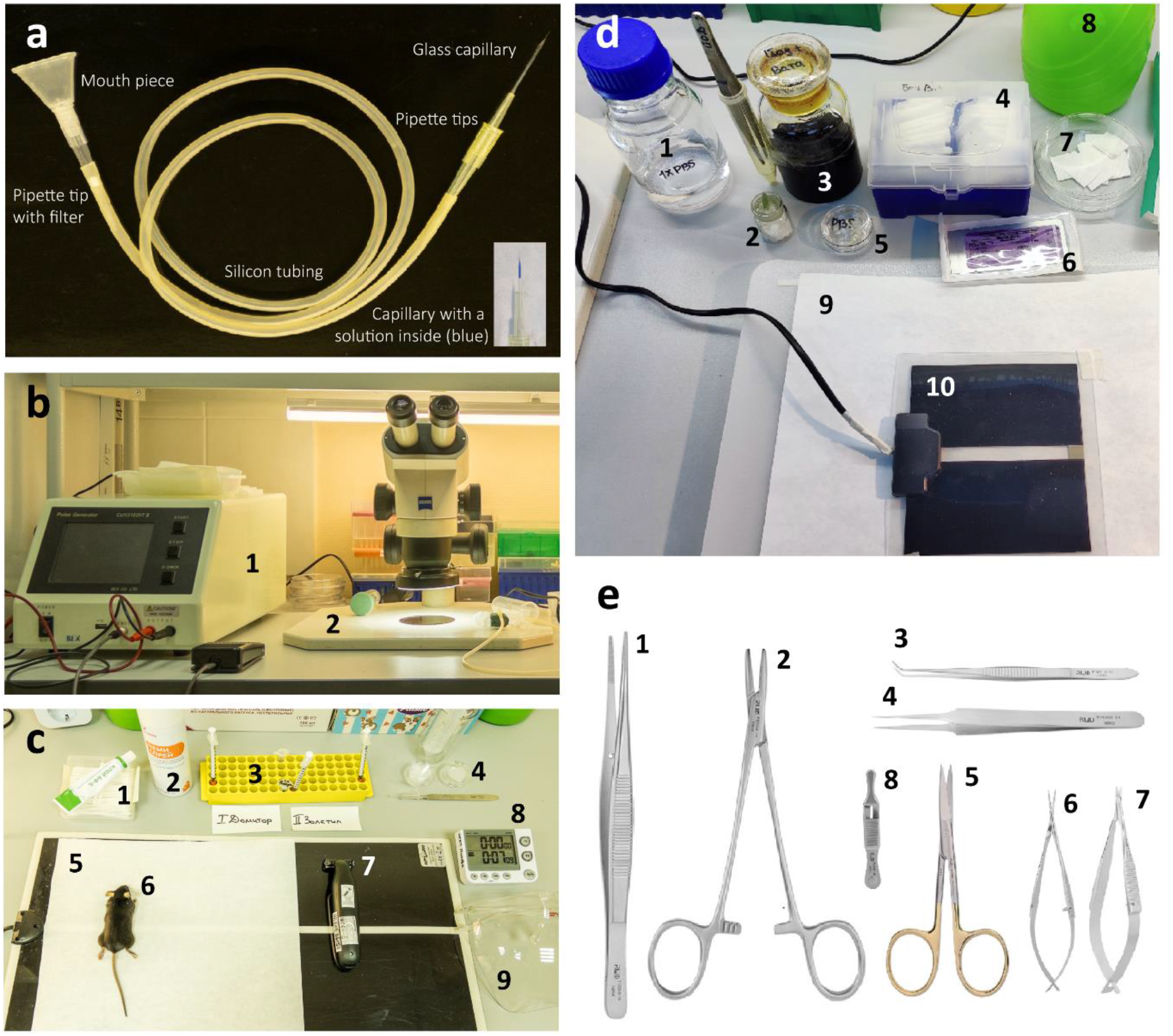
Pre-surgical preparation. **A. "Mouth pipette" system.** General view and enlarged image of a glass capillary with a solution inside (blue; bottom right). **B-D.** Workspace for intra-oviduct injection. **B.** Electroporation area. 1. Electroporator (BEX CO LTD, cat. #CUY21EDIT2); 2. Microscope (Zeiss, cat. #Stemi 2000). **C.** Pre-and post-surgical workspace. 1. Medical glue “BF-6”; 2. Chemi spray; 3. Insulin syringes with ready-made solutions; 4. Gel plates for eyes (3.5 cm Petri dishes with gel inside, scalpel (RWD, cat. #S31010-01, cat. #S32004-13) for cutting gel); 5. Thermal mat (28×53 cm) with filter paper (25×25 cm); 6. Anesthetized mouse; 7. Trimmer (hair clipper); 8. Stopwatch timer; 9. Protective shield. **D.** Workspace for surgical operation. 1. 1× PBS (filtered); 2. Covering wipes for the oviduct during electroporation (Kimtech Science, cat. #7552) (3×7 mm) in a small air-tight glass cup (sterilized); 3. Sterile cotton balls in an iodine alcohol solution (5%); 4. Covering wipes (10×10 cm) (Mastmed, Russia, cat. #613013) for the surgical field with 2×2 cm hole in the center of each wipe (sterilized); 5. 3.5 cm Petri dish with 1× PBS (filtered); 6. Surgical PGA absorbable suture with needle (Lintex, cat. #HR-20); 7. 10 cm Petri dish with filter wipes for removing blood and other liquids during surgery (1×1 cm) (sterilized); 8. Ethanol (70%) in a spray bottle (for disinfection); 9. Workspace for surgery ‒ the silicone mat (30×30 cm) with a sheet of filter paper (25×25 cm); 10. Thermal mat (14×15 cm). **E.** Surgical instruments. 1. Dressing forceps (RWD, cat. #F12016-15); 2. Needle holders (RWD, cat. #F31034-14); 3. Dressing forceps w/o serrations (RWD, cat. #F12013-10); 4. Micro forceps (RWD, cat. #F13002-11); 5. Scissors (round type) (RWD, cat. #S18004-09); 6. Scissors (triangular) (RWD, cat. #S11001-08); 7. Scissors (triangular) (RWD, cat. #S11035-08); 8. Clips (RWD, cat. #R33001-48).

#### Electroporation settings

In our work we use a square-wave pulse Electroporator (BEX CO LTD, cat. #CUY21EDIT2) as published previously (Ohtsuka *et al*., 2018). The following parameters must be set before starting: Pd A ‒ 100 mA; Pd on ‒ 5.00 ms; Pd off ‒ 50.00 ms; Pd cycle ‒ 3; Pd V ‒ 150 V; Decay ‒ 10%; Decay Type: Log; mode ‒ Square (mA) (+/−).

Pd ‒ driving pulses, which deliver DNA, RNA, etc. into the inside of the cell. Pd A ‒ maximum current. Pd on ‒ pulse length. Pd off ‒ pulse interval. Pd cycle ‒ number of pulses. Pd V ‒ voltage. Decay ‒ decay rate. Square (mA) mode ‒ constant current mode. Constant current mode provides stable current throughout the oviduct even though the manual grab is inconsistent over the trials. (+/−) ‒ the polarity of the square pulses alternately. The description of the modes is taken from the official webpage of the manufacturer (BEX CO LTD) (https://www.bexnet.co.jp/english/product/device/invivoinvitro/cuy21edit2.html; accessed on 5 September 2023).

### Surgical operation (*in utero* experiment)

#### Injection of anesthesia and analgesia

For anesthesia and analgesia, we used commercial drug “Domitor” (aqueous solution of medetomidine hydrochloride, 1 mg/ml) (for veterinary medicine) (Orion Pharma, Finland) and commercial drug “Zoletil 100” (aqueous solution medetomidine hydrochloride, 100 mg/ml) (for veterinary medicine) (Virbac, France). Anesthesia scheme (drug concentration) was carried out according to (Saydakova *et al*., 2023). Using this scheme allows the animal to be at rest for about 120-180 min. Detailed description of the procedure is the following:

1. With hands (in gloves) carefully remove the mouse by the tail from the cage and measure body weight, then return the animal to the cage.
2. Commercial drug “Domitor”: Open and put the date of opening on the vial, store the stock solution for no more than 3 months. Dilute the stock solution with saline 20 times (up to a concentration of 0.05 mg/ml). Dispense working solution into 1 ml aliquots and store at -20°C.
3. Commercial drug “Zoletil 100”: Prepare a stock solution (100 mg/ml) according to the manufacturer’s instructions. Dispense the stock solution into 50-100 µl aliquots in 1.5 ml tubes and store at -20°C. To prepare a working solution, dilute the stock solution 20 times with saline (up to a concentration of 5 mg/ml). The diluted drug should be stored for no more than 5 days at +4°С; do not freeze, as the effectiveness of the drug is reduced.
4. With hands (in gloves) carefully fix the mouse in the left hand: the thumb and forefinger fix the scuff of the animal, and the little finger presses the tail to the palm of your hand. Intraperitoneally inject diluted “Domitor” (concentration 0.05 mg/ml, which corresponds to 0.1 µg/µl) to mice in an amount of 10 µl per 1 g of animal body weight (i.e., 250 µl of the diluted solution per animal weighing 25 g) using insulin syringes. Return the animal to the cage.
5. After 10 min, intraperitoneally inject diluted “Zoletil 100” (concentration 5 mg/ml) into mice in an amount of 10 µl per 1 g of animal body weight (i.e., 250 µl of diluted solution per animal weighing 25 g) using insulin syringes.
6. After that, place the mouse on a thermal mat (35-37°С).
7. The depth of anesthesia is considered sufficient for a surgical operation if the anesthetized animal has no reflex reactions, the muscles are relaxed, and deep and rhythmic breathing is observed. Therefore, subsequent manipulations should be carried out 10 min after the injection of “Zoletil 100” for mice weighing ≥ 20-30 g or 15 min for animals weighing 35-40 g.
8. After entry into anesthesia, the eyes of the mouse should be covered with a plate of 1% agarose gel (prepared in water, room temperature after complete solidification) 3- 4 mm thick to prevent drying of the cornea (Figure S2.A). To prepare a gel plate of a convenient size, we use a Petri dish with a diameter of 3.5 cm, into which we pour 1% agarose gel. Next, the hardened gel is divided into two equal parts (semicircles) with a scalpel (RWD, cat. #S31010-01, S32004-13).

*Note: All following manipulations with animals should be carried out on a thermal mat (35-37°С)!*

#### Preparing an animal for surgery

9. Be sure to check the depth of anesthesia (the animal does not react to pinching fingers and tail). Then shave the operating area on the back with a trimmer. Perform the procedure in a protective shield or goggles to protect your face from animal hair.
10. Place the animal (back up) on a thermal mat (size 14×15 cm) with the heating temperature set to 35-37°C. Anchor the animal’s legs to the thermal mat with strips of painting tape (Figure S2.B).
11. Treat the mouse’s dorsal skin with ethanol (70%), wipe with a sterile cotton ball, then wipe with a sterile cotton ball soaked in iodine alcohol (5%) (Figure S2.B). Place covering material for the surgical field so that the 2×2 cm hole is centered over the incision site. Attach the covering material with the painting tape strips to the thermal mat (Figure S2.C).

#### Incision for intraperitoneal access to the oviduct

12. With one hand, lift the mouse’s dorsal skin with dressing forceps (RWD, cat. #F12016-15). Using scissors (RWD, cat. #S18004-09) with the other hand, along the spine in the region of 4-5 lumbar vertebrae make a sagittal (‘‘vertical’’) incision of about 1 cm (Figure S2.C-D). Do the work with the oviducts sequentially. First, perform all manipulations with the left oviduct. Soak filter wipe (size 1×1 cm) in 1× PBS (in sterile Petri dish, diameter 3.5 cm). Place the wipe to the right of the incision on the skin.
13. Place micro forceps (RWD, cat. #F13002-11) under the skin incision, shift to the left of the spine, lift the muscle tissue in the area of the physiological location of the left ovary (to the left of the vertebral muscle). Then using scissors make an abdominal incision no larger than 0.5 cm (Figure S2.E).
14. Then gently using micro forceps pull out the ovary, oviduct and part of the uterine horn and place this on a wet wipe (Figure S2.F). Attach bulldog clip (RWD, cat. #R33001-48) to the fat tissue above the ovary to prevent the organs from being pulled back into the abdominal cavity (Figure S2.G).

#### Entering the solution into the ampulla

15. Carefully place the mouse under the binocular microscope (without removing the animal from the heating mat) (Figure S2.H). Locate the oviduct ampulla (a portion of the oviduct showing oviductal expansion containing the fertilized zygotes) using a binocular.
16. With one hand, gently grab the oviduct near the ampulla with micro forceps so that it is convenient for you to insert the microcapillary with the solution into the ampulla. It is better to place the injection site of the solution closer to the beginning of the ampulla, retreating a few millimeters from the infundibulum. Please note that it is required to inject the solution in the direction of the "flow" of the zygotes from the infundibulum through the oviduct to the fallopian tube.
17. Using the "Mouth pipette" system with a ready-made microcapillary with a solution inside, blow the solution into the ampulla (Figure S2.I). The success of the injection can be understood by the fact that the injected solution (colored in blue) will be visualized inside the ampulla, and the ampulla will slightly increase in size (Figure S2.J).

#### Electroporation

18. Immediately after the solution enters the ampulla, electroporation procedure is performed. Wet a strip of sterile covering wipe (3×7 mm in size) in 1× PBS and cover the ampulla (Figure S2.K, arrow). Connect the tweezer-type electrode (BEX CO LTD, cat. #LF650P3) to the electroporator (BEX CO LTD, cat. #CUY21EDIT2). Then wet the ends of the tweezer in 1× PBS.
19. Next, gently (so as not to move the solution further down the oviduct) grab the ampulla with a tweezer-type electrode (Figure S2.L). It is important that the electrodes do not touch the tissue directly, but only through the strip of sterile covering wipe (3×7 mm in size). Perform electroporation with the settings set previously (described in the "Electroporation settings" section). An indicator of successful electroporation will be the appearance of air bubbles at the junction of the electrodes and ampulla.
20. Remove the strip of sterile covering wipe from the oviduct. While holding the fat tissue above the ovary, carefully detach the bulldog clip. Next, using micro tweezers, gently return the organs to their physiological location in the abdominal cavity.

**Figure S2.**
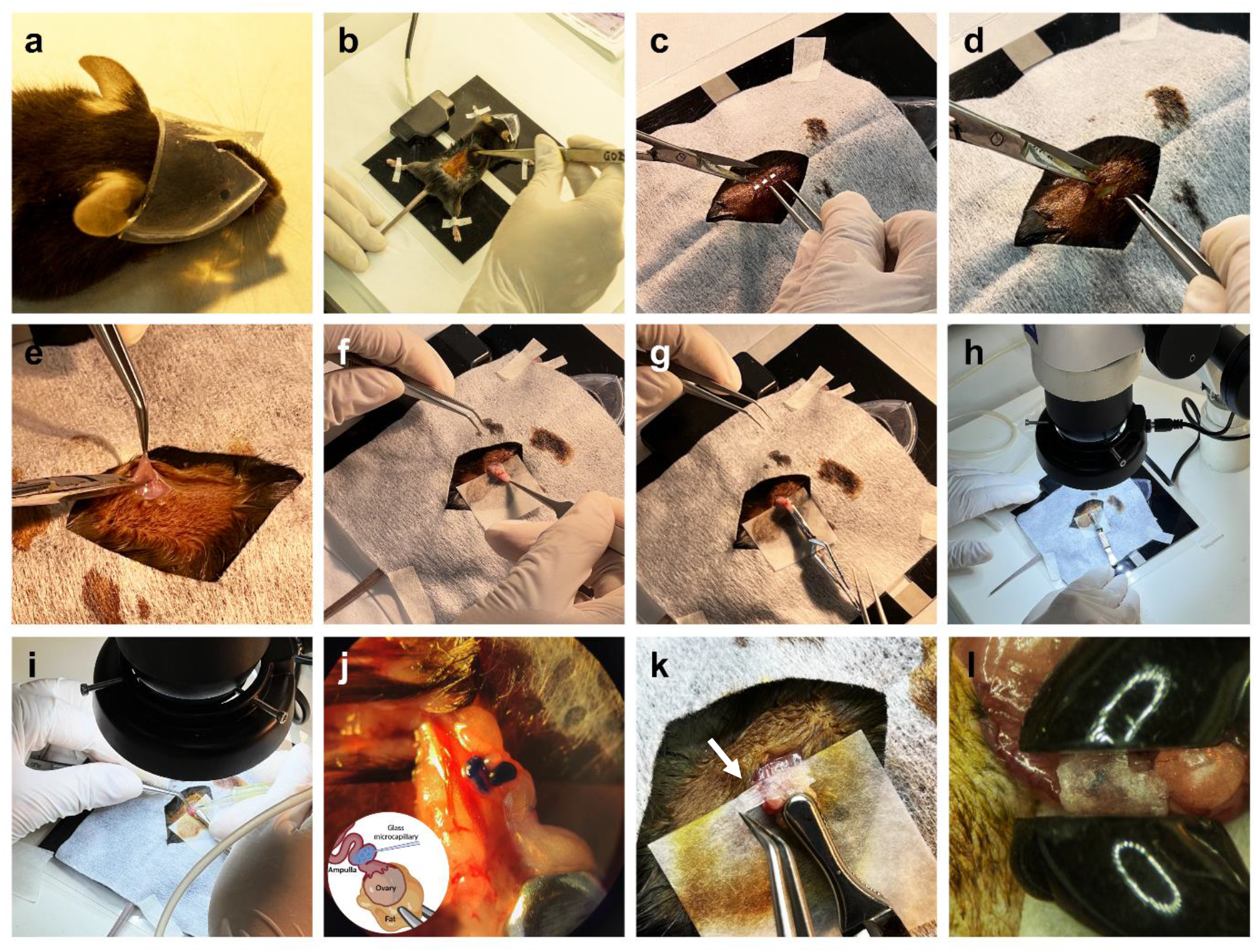
Mouse surgery (*in utero* experiment). **A-B. Preparing an animal for surgery. A.** Eyes of the mouse covered with a plate of 1% agarose gel. **B.** Animal placed (back up) on a thermal mat (35- 37°C), legs anchored to the thermal mat with strips of painting tape. Surgical area are treated with ethanol and iodine alcohol. **C-H**. Incision for intraperitoneal access to the oviduct. **C-D.** Place covering material for the surgical area. Made a sagittal incision of about 1 cm (dotted line). **E.** Made an abdominal incision no larger than 0.5 cm. **F.** Pulled out the ovary, oviduct and part of the uterine horn and placed on a wet wipe. **G.** Attached clip to the fat tissue above the ovary. **H.** Placed mouse under the binocular microscope. **I-L**. Entering the solution into the ampulla and electroporation. **I-L.** Injection solution into the ampulla using the "Mouth pipette" system. **K.** Covered with wet wipe (3×7 mm) ampulla (arrow). **L.** Grabbed ampulla with a tweezer-type electrode.

Perform the manipulations to introduce the solution into the ampulla of the right oviduct as described above (par. 13-20). For convenience (if you are right-handed), turn the thermal mat with the mouse 180 degrees (so that the mouse is facing you).

#### Suturation

21. When the right oviduct is completed, add 100 µl of 1× PBS into the peritoneal cavity and suture both muscle incisions and one in the skin with surgical PGA absorbable suture with needle suture and needle (Lintex, cat. #HR-20). Use an interrupted vertical suture, which is most often used to close postoperative wounds. Conjunction of the tissue edges should be done without tension to avoid tissue damage.
22. The needle should be inserted facing the operator ‒ from the far edge and come out from the near. Fix the far edge of the wound with forceps and carefully insert the needle at a distance of 0.15-0.2 mm from the edge, then pass the tip and part of the needle body through the tissue, fixing the edge of the wound with the forceps.
23. Then take the near edge of the wound and pass the tip and body of the needle through the tissue. After this, the needle must be intercepted with needle holders (RWD, cat. #F31034-14) and further moved pulling the thread. Tighten the seam. Make a square knot. Repeat and make 2-3 stitches depending on the size of the wound (Figure S3.A-C).

#### Post-surgical procedures

24. After the mouse’s dorsal skin has been sutured, post-surgical care must be taken. Remove the covering material from the animal, and use Chemi spray or analogous anti-inflammatory and antibacterial drug to treat the wound. After the solution has dried, close the seam with medical glue “BF-6” or similar (Figure S3.D). It will prevent the mouse from removing the suture until the wound heals.
25. To relieve pain syndrome after the animal comes out of anesthesia, we use the drug "Rimadyl". It must first be diluted, since the initial concentration (5%) is applied for larger animals (cats, dogs). Prepare the working solution as follows: dilute the stock solution (5%) 20 times (50 µl drug “Rimadyl” + 950 µl normal saline). Next, dilute the resulting solution by another 1.5 times (add 400 µl solution to 300 µl normal saline). Store the final solution at +4C for no more than 2 months. Inject the working solution subcutaneously in the scuff by 100 µl per mouse weighing 30 g.
26. After the end of all procedures, it is necessary to replace the gel plate to protect the eyes of the mouse with a new one. Then put the animal in a clean cage placed on a thermal mat size 28×53 cm (set the heating temperature to 35-37°С). The cage with the animal is kept on a warm mat until the animal comes out of anesthesia. There is no need to cover the animal, since the covering material can stick to the medical glue “BF-6” and this will create discomfort for the animal in the future.

If necessary, perform the above manipulations with more animals. Important: before starting work with the next animal, it is necessary to treat all instruments with a disinfectant, for example “Kombidez” (KiiltoClean, Russia). To do this, we pour approximately 25-30 ml of solution into a 50 ml falcon and dip the working surface of the instruments into it several times. We recommend working in pairs, as this will save time. After electroporation of both oviducts, the animal is transferred to the second operator for further suturation and treatment of wounds (par. 21-26). The first operator starts the manipulations (par. 19-20) with a new animal.

### Post-surgical care

Within 4-5 days after the operation, each animal is checked for the integrity of the suture and wound healing. Be sure to treat the sutures with Chemi spray to accelerate the healing of the suture, and this also prevents the animals from interfering with wound healing (Figure S2.E).

**Figure S3.**
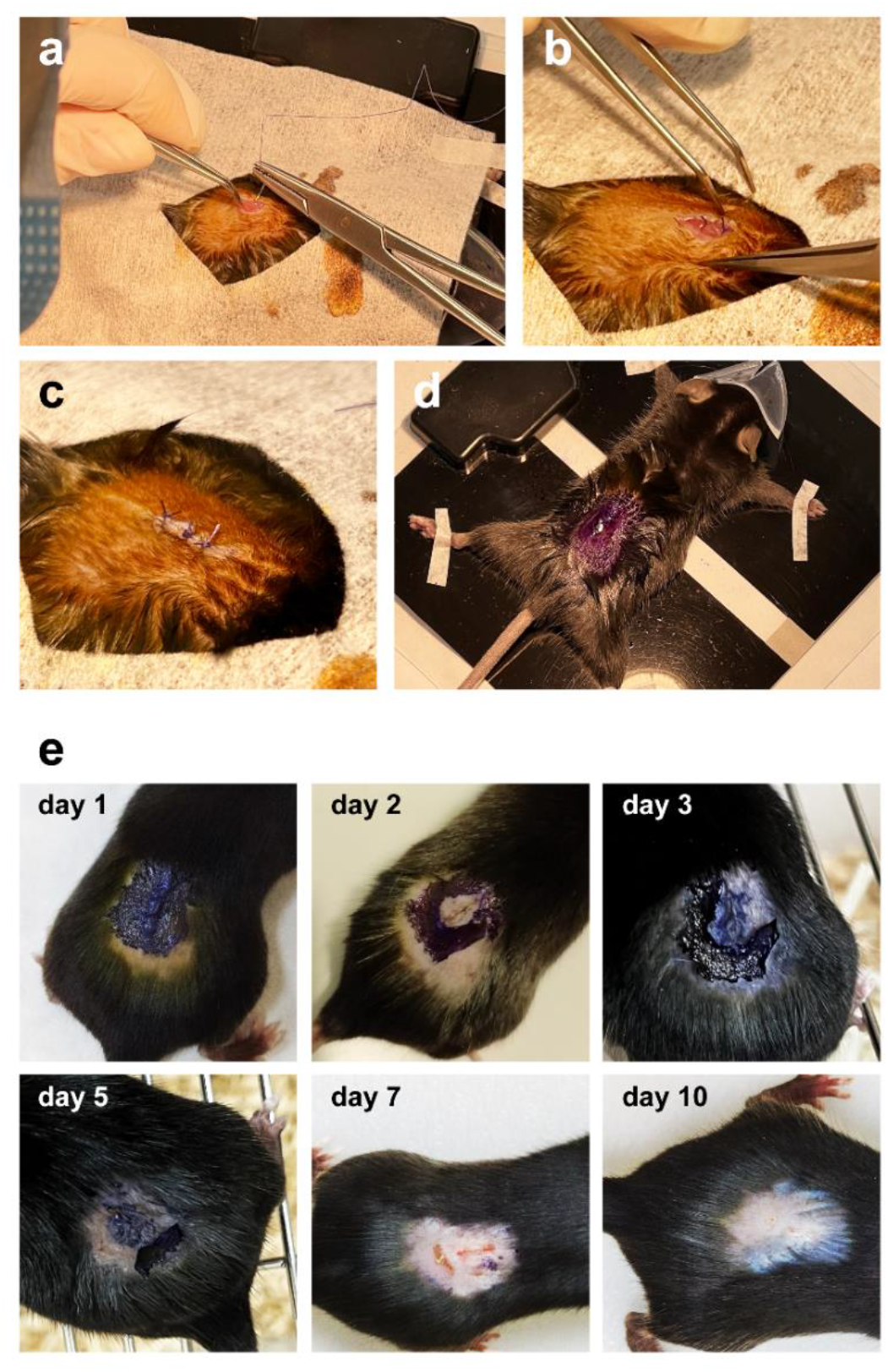
A-D. Suturation and post-surgical procedures. **A-B.** Suturing on muscle incisions (interrupted vertical suture). **C.** Suturing on skin incision (interrupted vertical suture). **D.** Reduce inflammation procedures with Chemi spray and medical glue “BF-6” to protect the suture from mouse’s removing. **E.** Wound healing after surgery: day 1-10.

### Testing electroporation procedure in embryos

To determine the damage of the electroporation procedure and the potential embryotoxicity from *IL10* construct, we decided to make two groups: control and experimental. All manipulations were performed on anesthetized CD-1 pregnant females (0.7 d.p.c.) without hormonal stimulation (all from the points “pre-surgical preparation” and “surgical operation”, with the exception of “suturation” and further procedures). In the experimental group, the *IL10* sgRNA (30 µM) in Opti-MEM I Reduced Serum Medium (ThermoFisher Scientific, cat. #31985062) was introduced into the ampulla and then electroporated. In control group the mix of 1× PBS (instead of *IL10* sgRNA) and Opti-MEM I Reduced Serum Medium was also introduced into the ampulla and the electroporation procedure was carried out on the same settings.

Next, in both groups, zygotes were washed out of the ampulla. To do this, the oviduct isolated from the female and a solution of KSOM Embryo Medium (Sigma, cat. #MR-101-D) with hyaluronidase (Roanal, cat. #08091) (10 mg/ml in KSOM Embryo Medium) using the “Mouth pipette” system was injected into oviduct until all zygotes were completely washed out. Next, the zygotes were placed into fresh portion of KSOM Medium onto a culture plate covered with mineral oil (Sigma-Aldrich, cat. #M8410-100ML). Then the zygotes were incubated at a temperature of 37°C in 5% CO_2_ for 24 h, after which the number of dividing cells was counted.

To confirm electroporation efficiency, Fluorescein isothiocyanate-dextran (FITC-dextran) (Sigma-Aldrich, cat. #FD4-1G) was diluted in concentration of 10 µg/µl in Opti-MEM medium, introduced into the ampulla of CD-1 mice in volume 1.5 µl/oviduct followed by electroporation. Zygotes were washed out and analyzed using fluorescent microscope.

### Offspring genotyping

On days 21-22 after the i-GONAD procedure, female mice give birth to pups. It is necessary to check the nest for the presence of dead pups, because C57BL/6 female mice can cannibalize their offspring. The dead pups must be removed from the cage, and a tissue sample should be kept for genotyping (the tail is most convenient). The surviving pups are left until weaning at the age of 4-5 weeks. Further, the pups must be separated from the mother in fresh cages (same-sex siblings are kept together).

#### Extraction of genomic DNA from tails and ears (chloroform extraction and precipitation)

1. Before taking the material, autoclave and dry the 1.5 ml tubes. Collect small (about 5 mm) pieces of tissue (ears or tails) into test tubes, label the tubes. Store samples at - 20°C or -70°C until DNA is isolated.
2. Prepare buffer “A” (50 ml volume): TrisHCl (1M, pH=8,0) (2.5 ml, final concentration 50 mM), NaCl (5M) (1 ml, final concentration 100 mM), EDTA (0.5M, pH=8.0) (2.5 ml, final concentration 25 mM); mQ H_2_O (44 ml). Store buffer “A” at room temperature.
3. Prepare lysis buffer (10 ml volume): 10% SDS (500 µl, final concentration 0.5%), buffer “A” (9.5 ml). Do not store lysis buffer more than 1 day after preparation.
4. Add 250 µl of lysis buffer and 2 µl of Proteinase K (20 mg/ml; Sibenzyme, cat. #E347) per sample. Incubate samples at 56°C overnight.
5. Homogenize the samples, collect drops using a table-top centrifuge. Add 105 µl of KAc (5M) to the samples, mix thoroughly, collect drops using a table-top centrifuge (at this stage, the proteins remaining in the samples precipitate, the solution becomes white). Add 375 µl of chloroform, mix. *ATTENTION: Chloroform extraction should be performed in a fume hood!*
6. Centrifuge at 12000 rcf for 10 min at 20°C. After centrifugation, the sample is divided into fractions: the lower one is chloroform; interphase - proteins, debris; the upper one is the aqueous phase containing DNA. Prepare clean 1.5 ml tubes, sign according to the samples to be isolated. Carefully (without touching the interphase and chloroform!) take 180 µl of the aqueous phase into one clean tube. *ATTENTION: After extraction, drain the rest of the contents of the test tubes into the “Organic Waste”, leave open test tubes and plastic to ventilate under the hood!*
7. To 180 µl of the aqueous phase add 540 µl of 96% ethanol, mix thoroughly and incubate for 60 min at -20°C. Centrifuge at 12000 rpm for 15 min at 20°C. After centrifugation, a white precipitate of DNA is observed at the bottom of the tube. Carefully remove the supernatant without touching the DNA pellet. Add 700 µl of 70% ethanol, mix. Centrifuge at 12000 rpm for 5 min at 20°C.
8. Carefully remove the supernatant without disturbing the DNA pellet. Dry the precipitate. *Note: It is necessary that the alcohol evaporates completely!* Add 50-100 µl mQ H_2_O, mix thoroughly. Determine the DNA concentration and check the purity of the samples using NanoDrop (blank ‒ mQ H_2_O).

#### Preparing PCR samples

For preparing PCR samples BioMaster HS-Taq PCR-Color (2×) (Biolabmix, cat. #MHC010-200) was used according to the manufacturer’s protocol. Per reaction was taken 100 ng of DNA sample. DNA samples from intact C57BL/6 animals were used as a positive control and mQ H_2_O as a negative control. The following primers were used for PCR: IL10_ex1_gDNA_F / IL10_ex1_gDNA_R (259 bp PCR product size), IL10_ex2_gDNA_F / IL10_ex2_gDNA_R (371 bp PCR product size), IL10_ex2_gDNA_F1 (5’ AAACTCTCCTTTCCACAGTTGC 3’) / IL10_ex2_gDNA_R (475 bp PCR product size).

After the PCR is completed, the samples are loaded onto a 2% agarose gel prepared on 0.5× TBE (Tris/Borate/EDTA) buffer. Samples of the correct size (Table S3) were cut out of the gel using a transilluminator. PCR products were isolated from the gel using a DNA and RNA agarose gels isolation kit (Biolabmix, cat. #N-Gel-250) according to the manufacturer’s protocol.

#### Sequencing (according to Sanger)

The DNA nucleotide sequence was determined using the BigDye Terminator v3.1 cycle sequencing kit (ThermoScientific, cat. #4337455) according to the manufacturer’s protocol. The PCR product after isolation from agarose gel was used as a template. Primers IL10_ex1_gDNA_F, IL10_ex2_gDNA_F, and IL10_ex2_gDNA_F1 were used for sequencing. Sequencing was performed using an Applied Biosystems 3500 Genetic Analyzer capillary sequencer (Applied Biosystems, USA).

## TROUBLESHOOTING

In the first experiment, we performed the i-GONAD procedure according to Ohtsuka et al (Ohtsuka *et al*., 2018), but instead of isoflurane anesthesia, we used the “Domitor + Zoletil” (Orion Pharma, Finland; Virbac, France) anesthesia scheme described in the methods above. However, we did not obtain either offspring or pregnant females. We do not attribute this failure to the anesthesia used, since the “Domitor + Zoletil” anesthesia scheme is used by many scientific groups. (Cagle *et al*., 2017; Limprasutr *et al*., 2021; L.W. Hall, 2002) However, as a result of the first experiment, a number of problems were noted: 1) mouse fur soiled the wound; 2) test mice quickly removed the sutures; 3) cornea dried during surgery despite we applied eye cream as suggested in published protocols.

Thus, starting the second experiment, we addressed the following issues:

1. We shaved fur in the surgical area to avoid it getting inside the wound both when cutting the skin and when applying sutures. We noted that shaving the fur around the skin incision site had a positive effect on wound healing.
2. To reduce the risk of sutures being removed by mice in the first hours after surgery, we decided to use a special collar for rodents (for mice, neck diameter 1.25’, KVP, cat. #RCM-1’) in the second attempt. The use of a collar for rodents had a negative impact on the animals, as the mice showed obvious stress due to the inability to perform grooming. Taking into account the experience of the second experiment, it was decided not to use a collar for rodents in the future. Instead, we applied medical glue “BF-6” (Verteks, Russia) to protect the seam from being removed by a mouse. To improve wound healing, we used Chemi spray, an anti-inflammatory and antibacterial drug (Industrial Veternaria S.A. Invesa, Spain). Since mice can remove stitches due to pain, we used the analgesic and anti-inflammatory drug “Rimadyl” (Zoetis, Brazil) to reduce pain after anesthesia
3. During anesthesia of animals, it is necessary to moisture the cornea of the eye, since the corneal reflex is absent in animals under anesthesia. To eliminate dry eyes, we used ophthalmic eye hydrogel “Vidisik”, 0.2% (Dr. Gerhard Mann Chemical-Pharmaceutical Company GmbH, Germany). It partially helped to prevent the cornea from drying, however, this hydrogel dries very quickly and damages the cornea. Therefore, we decided to use agarose gel pads to protect eyes from damage during the surgery, which worked very well for us.

